# Statistical method accounts for microscopic electric field distortions around neurons when simulating activation thresholds

**DOI:** 10.1101/2024.10.25.619982

**Authors:** Konstantin Weise, Sergey N. Makaroff, Ole Numssen, Marom Bikson, Thomas R. Knösche

## Abstract

**Introduction:** Notwithstanding advances in computational models of neuromodulation, there are mismatches between simulated and experimental activation thresholds. Transcranial Magnetic Stimulation (TMS) of the primary motor cortex generates motor evoked potentials (MEPs). At the threshold of MEP generation, whole-head models predict macroscopic (at millimeter scale) electric fields (50-70 V/m) which are considerably below conventionally simulated cortical neuron thresholds (175-350 V/m).

**Methods:** We hypothesize that this apparent contradiction is in part a consequence of electrical field warping by brain microstructure. Classical neuronal models ignore the physical presence of neighboring neurons and microstructure and assume that the macroscopic field directly acts on the neurons. In previous work, we performed advanced numerical calculations considering realistic microscopic compartments (e.g., cells, blood vessels), resulting in locally inhomogeneous (micrometer scale) electric field and altered neuronal activation thresholds. Here we combine detailed neural threshold simulations under homogeneous field assumptions with microscopic field calculations, leveraging a novel statistical approach.

**Results:** We show that, provided brain-region specific microstructure metrics, a single statistically derived scaling factor between microscopic and macroscopic electric fields can be applied in predicting neuronal thresholds. For the cortical sample considered, the statistical methods match TMS experimental thresholds.

**Conclusions:** Our approach can be broadly applied to neuromodulation models, where fully coupled microstructure scale simulations may not be computationally tractable.

## 1. Introduction

Transcranial magnetic stimulation (TMS) of the primary motor cortex (M1) causes peripheral muscle activation, reflected by motor evoked potentials (MEP) recorded from surface electrodes over respective muscles. Such experiments are valuable for studying the motor system and its pathologies (e.g., Di Lazzaro & Ziemann, 2013), and underpin individual dosing of repetitive TMS (rTMS) therapies, such as for depression (Rossi et al., 2021). The amplitude of the MEP scales with the TMS device output and, more directly, with the electric field the relevant neurons are exposed to. Modeling the relation between stimulation intensity and cortical responses underpins explaining TMS and rTMS outcomes.

Numerical field modeling in conjunction with a non-linear regression approach has enabled localization of activated neuronal populations and the derivation of input-output (IO) curves that map TMS induced electric field strengths to the MEP amplitudes (Weise et al., 2020; Numssen et al., 2021; Weise et al., 2023a). With conventional biphasic pulses these sigmoidal IO dose responses have a half-maximum (50% of peak MEP response) at ∼50-70 V/m (Numssen et al., 2021). However, explicit simulations of cortical L3/4 neurons predict higher thresholds of ∼260 V/m for the same TMS waveform (Weise et al., 2023b) and 175-350 V/m for monophasic pulses (Aberra et al., 2020; Weise et al., 2023b). We hypothesize that this mismatch is a consequence of conventional numerical field models ignoring the presence of microscopic structures (cell membranes, blood vessels).

In calculation of electric fields produced during neuromodulation (TMS), classical models assume macroscopically (mm scale) homogenous tissue. At the microscopic (µm) scale, however, the conductivity is highly inhomogeneous given the low conductivity of cell membranes and vasculature. Recently, this effect has been investigated in detail by Qi et al. (2025) using a high-resolution boundary elements model of a sub-volume (250×140×90 μm) of the L2/3 P36 mouse primary visual cortex with detailed segmentation of microscopic compartments, taking into account neuronal and glial membranes as well as blood vessels (Turner et al. 2022; MICrONs Consortium 2021), and comprising ∼0.5 billion facets in total.

Adapted from (Qi et al., 2025), Figure 1 shows how low conducting barriers cause charge accumulation and Fig. 2 demonstrates how this changes the effective electric field at the cell membranes. This electric field inhomogeneity may explain the apparent discrepancy between the activation thresholds found in microscopic simulations to those obtained when relating experimentally observed MEPs to macroscopic electric field simulations of TMS. The goal of this work is to test this hypothesis.

**Fig. 1:**
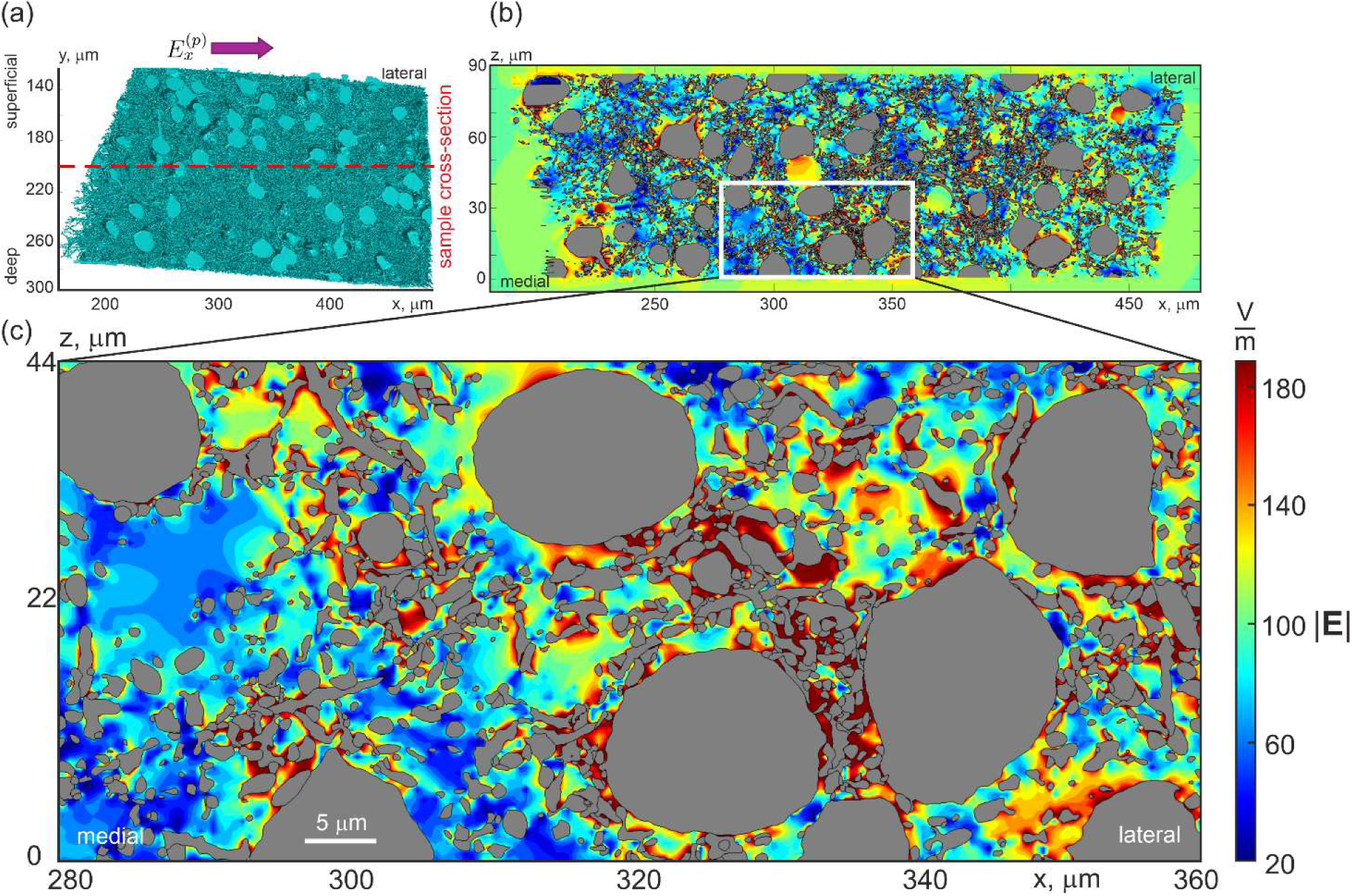
Distribution of the extracellular electric field magnitude (V/m) inside the sample when a uniform impressed electric field of 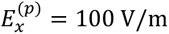 is applied in the medial-lateral direction (from left to right in the (a)), showing how low conducting barriers (e.g., cell membranes) cause charge accumulation and associated field distortion in (b) and (c). The intracellular space (grey) is not included in the computations. Tissue segmentation and electric field computations are from Qi et al. (2025).

**Fig. 2.**
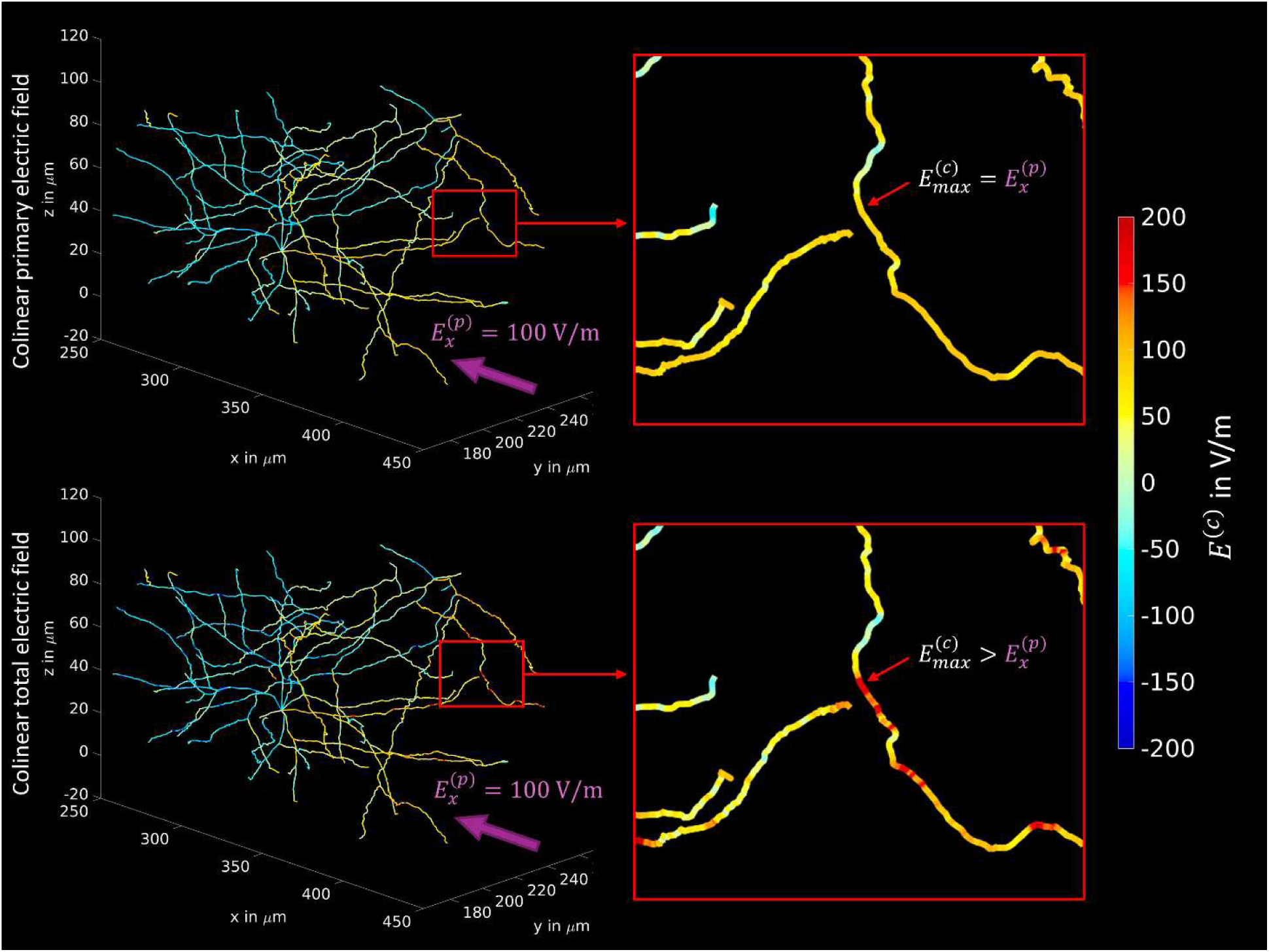
The influence of the accumulation of charges on the membranes onto the collinear electric field at the centerlines of neuronal processes for an example neuron. Top: A uniform impressed electric field is applied along the *x*-axis (from dorsal to ventral) with 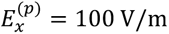. The influence of the charges is not considered and the impressed field is directly projected to the centerlines. Hence, the maximum achievable collinear field (for an optimally oriented neuronal segment) at the neurons 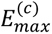 is equal to the impressed field. Bottom: The same uniform impressed field is applied, but realistic neuronal (sub-)compartments are included into the voltage simulation and the impressed field is distorted by the field of the induced charges. In consequence, the maximum achievable collinear field 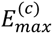 can be larger than the impressed field 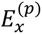.

To this end, we use the electric field simulations performed by Qi et al. (2025) to determine the microscopic fields at the axons that correspond to a given macroscopic field. As prior simulations (Aberra et al. (2020); Weise et al. (2023b)) have revealed that the axonal terminals have the lowest threshold, we consider field differences between both approaches at axon terminals to estimate the recruitment rate as a function of the macroscopic field. If the microscopic electric fields from Qi et al. (2025) elicit action potentials at the axon terminals for macroscopic field strengths of 50-70 V/m (Numssen et al., 2021), then microstructure-dependent electric field warping does indeed account for the aforementioned activation threshold discrepancy seen in between conventional (macroscopic) and explicit (microscopic) modeling.

Since realistic microscopic simulations are computationally expensive, it can be impractical to directly replace the macroscopic field calculation methods with microscopic simulations in routine analysis. For this reason, the second aim of this study is to provide an easy-to-compute method that corrects the discrepancy between activation thresholds (or recruitment rates) based on microscopic and macroscopic fields. Our method offers a principled way to create, for any neuromodulation technology and any cortical tissue with available microanatomical representation, a lookup table that maps macroscopic field strength and orientations to recruitment rates.

## 2. Materials and Methods

### 2.1 Modeling framework

The present multiscale modeling framework includes three distinct yet interconnected steps, each utilizing a different numerical method:

i. **Biophysical computations**: Activation thresholds for rodent neurons with neuron morphologies from Markram et al. (2015) are computed under external brain stimulation. These computations are performed using the 1D cable equation with appropriate channel parameters.
ii. **Novel statistical model**: Essentially random microscopic field (extracellular potential) fluctuations in the cortical gray matter are statistically incorporated into the standard multiscale brain stimulation pipeline (Aberra et al., 2020; Weise et al., 2023b), which previously accounted only for macroscopic deterministic (smooth) brain stimulation fields.
iii. **Representative set of microscopic field/potential fluctuations**: A realistic set of extracellular microscopic field/potential fluctuations is derived from Qi et al. (2025). This set is used to initialize the parameters of the proposed statistical model.

Below, we describe each modeling step in detail.

### 2.2 Threshold computations in morphologically realistic neuron models

In extension of an earlier study by Aberra et al. (2020), Weise et al. (2023b) computed the thresholds of different neuronal populations with respect to the electric field at the membrane. Detailed models of a large number of neuronal morphologies, taken from the Blue Brain Project (Markram et al. 2015), were used to account for the natural variability of neurons of different types. These types include layer 2 and 3 pyramidal cells (L2/3 PC), layer 4 small, nested, and large basket cells (L4 S/N/LBC), as well as layer 5 pyramidal cells (L5 PC) from mouse cortex. The neural morphologies were scaled up to human dimensions. In those simulations, we assumed a homogeneous or linearly changing field across the neuron, thus neglecting any effects of the presence of the neurons themselves and other structures. This essentially means treating microscopic and macroscopic fields (see Introduction) as equal. The firing thresholds were determined independently for each neuron by applying external electric fields with different angles with respect to and different gradients along the somato-dendritic axis. Importantly, it turned out that the initial generation of action potentials (i.e., the excitement of the respective neuron) almost exclusively occurred at one of the axonal terminals. The excitation then spread over the entire axonal arbor and activated all other synapses. We utilize these results in the present study.

### 2.3 Alteration of neuronal recruitment rate due to microscopic field perturbations

To allow drawing general conclusions regarding the relevance of differences between microscopic and macroscopic electric fields on the apparent excitation threshold of neurons with respect to the macroscopic field, the problem must be treated statistically. The ratio between microscopic and macroscopic electric fields at an arbitrary location *r* on the (axonal) cell membrane 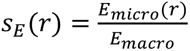 can be considered a random variable with a probability density *p*(*s_E_*). Estimating this distribution requires separate electric field simulations of (homogeneous) macroscopic and (inhomogeneous) microscopic electric fields in a sample of neural tissue, which are described in the next section. This provides the essential means to adapt the previously determined thresholds and recruitment rates of Aberra et al. (2020) and Weise et al. (2023b) with respect to microscopic electric fields. The corrected recruitment rate *r_s_* of a *single* axon terminal is then given as a function of the external macroscopic electric field *E_macro_* by:

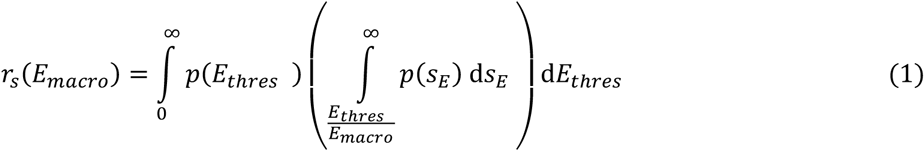

where *E_thres_* and *p*(*E_thres_*) are the threshold field strength and its probability density, respectively, calculated over different samples of neurons (within a particular neuron type) with externally applied electric fields from, e.g., Weise et al. (2023b). The inner integral 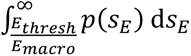 describes the probability that, at any arbitrary axonal terminal, the (relative) microscopic field *s_E_ = E_micro_/E_macro_* is above a given (relative) threshold *E_thres_/E_macro_*. The outer integral then sums that value over all possible threshold values, weighted by their probabilities, thus yielding the final activation probability (recruitment rate) of axonal terminals.

The fact that excitation of *any* axonal terminal of a given neuron eventually excites the entire axonal arbor with all its terminals (see previous section) leads to a statistical problem that depends on the average number of terminals *N* per neuron in the respective cell population. Accordingly, the recruitment rate of a population of neurons *r_n_*(*N_macro_*) can be determined from the recruitment rate from a single axon segment *r_s_*(*E_macro_*) from eq. (1) as follows:

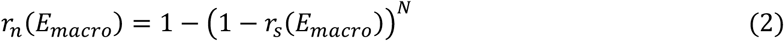

### 2.4 Computing the microscopic extracellular electric field in cortical cube sample

To determine the statistics for the scaling factor between microscopic and macroscopic electric fields *p*(*s_E_*), we utilized extensive electromagnetic numerical simulations of a neural tissue sample performed with boundary element fast multipole method (BEM-FMM) (Makarov et al 2018, Noetscher et al., 2023). The details and results of these computations are reported elsewhere (Qi et al., 2025). In short, the simulations were based on a 250×140×90 μm cube of neural tissue from mouse primary visual cortex, obtained from electron microscopic images with a resolution of 3.6×3.6×40 nm (the “pinky” dataset of IARPA Phase I). The reconstruction comprises triangulated surfaces (membranes) of various cellular structures, including pyramidal and non-pyramidal neurons, astrocytes, microglia, oligodendrocytes and precursors, pericytes, vasculature, nuclei, mitochondria, etc. (Turner et al. 2022; MICrONs Consortium 2021). The surface resolution (or average computational mesh size) was set at 100 nm to account for microscopic closely spaced objects.

The computation of the extracellular electric field is based on the assumption that membranes are non-conducting at the end of an initial polarization period, allowing to solve the extracellular field problem as a decoupled problem, using Neumann boundary conditions at the outer surfaces of the membranes (for details, see Noetscher et al., 2023; Makaroff et al., 2023; Qi et al., 2025). This implies that intracellular fields are not computed. Intracellular space is considered isopotential in the approximation of initial polarization, and intracellular structures, like mitochondria and nuclei, thus have no influence.

For the numerical treatment of the problem, the BEM-FMM method (Makaroff et al. 2018; Makaroff et al., 2023; Noetscher et al., 2023) was used. This method computes the electric charges deposited (or induced) on membrane surfaces in response to an impressed electric field. These charges (more precisely, the surface charge density) are responsible for the secondary electric field, which serves as the source of microscopic electric field and extracellular potential perturbations. The secondary extracellular electric field along its skeleton centerlines was forced to have the mean value zero to assure that the total extracellular electric field for every neuron within the sample has exactly the same mean value as in the homogeneous space.

According to classical quasi-static conduction theory (Wang et al., 2024; Jackson 1998, Balanis 2023, Stratton 2006), the resulting field perturbations are relatively strong (on the order of the impressed field itself), but highly localized. These perturbations typically have a spatial extent similar to the size of the perturber itself, such as the cross-section of a soma, dendrite, or axon. However, perturbations of the same polarity can potentially form large clusters, for example, in regions with several closely spaced somas. The size of such clusters may approach the axonal space constant, thereby facilitating easier excitation or inhibition of action potentials.

The BEM-FMM approach was specifically adapted to a large neuronal ensemble with several hundreds of closely spaced neurons and with the overall size of 0.5-1 billion facets, using a nested iterative algorithm (see Qi et al., 2025). Its idea is to compute charge deposition for each neuron (a membrane-based surface mesh with ca 1-2 million facets) separately, then update this charge deposition by the effect of all other neurons, and finally repeat this process iteratively until a proper convergence criterion is met. While the electromagnetic computational effort has been quite extensive and took over half a year, its accuracy has been verified by excellent self-convergence. Since the threshold computations by Weise et al. (2023b) and Aberra et al. (2020) (see Section 2.1) are based on one-dimensional cable equations, we need the extracellular potential or the collinear extracellular electric field at the centerlines of the neuronal processes. In the computations by Qi et al. (2025), however, neurons are intrinsically modeled as three-dimensional objects. Therefore, for performing the biophysical modeling, we extracted the average solution over the cross-sections of the processes (dendrites, axons, soma) as described in Qi et al. (2025).

## 3. Results

### 3.2 Microscopic electric field variations

Figure 2 illustrates, for an example neuron, the impact of the presence of neurons and other structures on the local collinear electric field at the membranes. In this particular case, we observe an elevation of the field maximum by more than a factor of 2. For better visualization, the values were limited to ±200 V/m, as extreme values can be up to a factor of 10. However, due to the complex morphology and mutual influences, the locations of field increases and attenuations cannot be assessed deterministically on a larger scale. This motivates a statistical approach to the problem, as described in Section 2.2, in order to account for the field variability of the microscopic electric field when calculating the neuronal thresholds and to correct the values determined by Aberra et al. (2020) and Weise et al. (2023b).

### 3.2 Influence of microscopic electric field variations on neural excitability

In total, the activation functions of ∼1.4 ⋅ 10^6^ axon segments were extracted from the cube sample. The histogram and the probability density function are shown in Fig. 3. According to that distribution, the median of the scaling factor is 1.06 and the probability that the electric field at an axon segment is higher than macroscopically assumed is 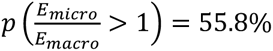. The distribution also shows that axon segments can be exposed to field strengths exceeding macroscopic electric field approximations by a factor of 5 and more. However, also the opposite can be the case, as axon segments may be exposed to very low field strengths, too.

**Fig. 3:**
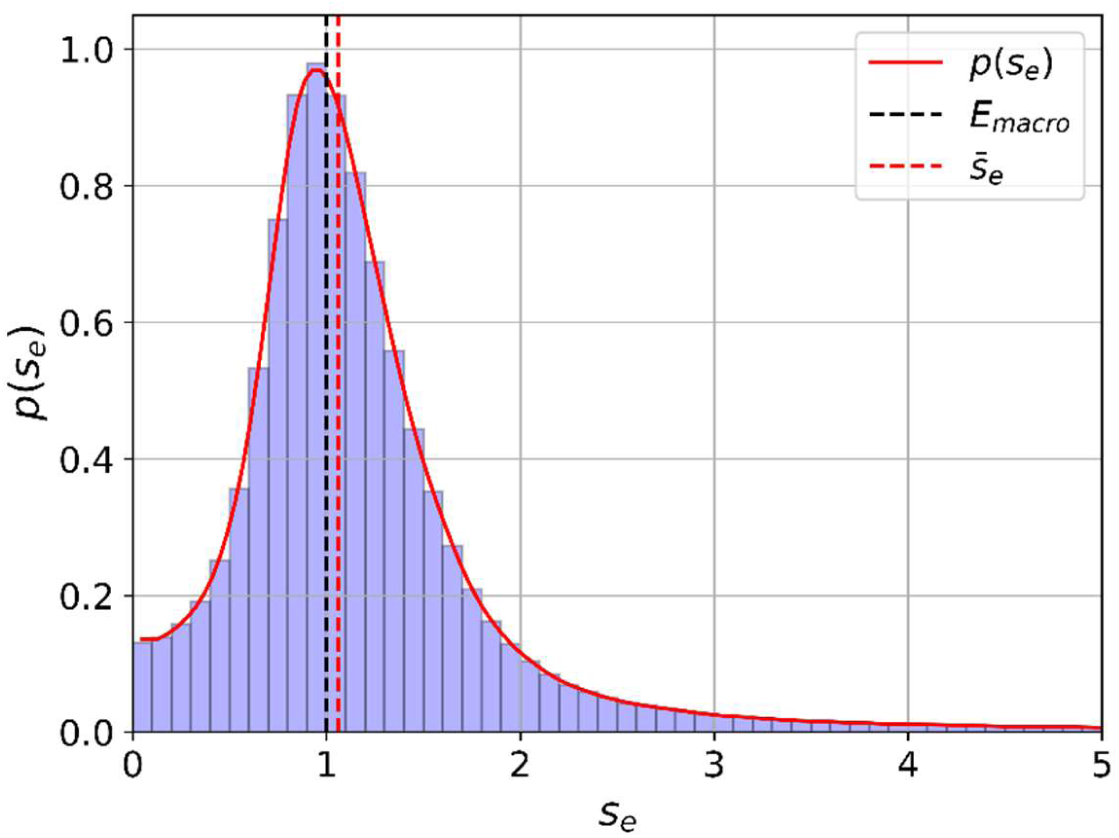
Histogram and probability density *p*(*s_e_*) of the electric field scaling factor 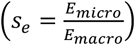 between microscopic and macroscopic electric fields (black dashed line). The median of the scaling factor is 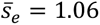 (red dashed line).

The probability density function of the electric field scaling factor (Fig. 3), in conjunction with the results from Weise et al. (2023b), allows for a realistic estimation of the neuronal recruitment rates with respect to the macroscopic field, using eqs. (1) and (2). Fig. 4 shows an example of a corrected recruitment rate taking into account microscopic electric field variations for the case of L2/3 PC from Weise et al. (2023b), where it was assumed that the macroscopic electric field is homogeneous and points along the somato-dendritic axis towards the soma. We assumed an average number of terminals per neuron of N=35 (STD=13.7), informed by the L2/3 population used by Weise et al. (2023b). The grey shaded lines are determined after sampling the number N of terminals per neuron from a normal distribution with the given mean and standard deviation to illustrate the expected variability of the recruitment curves. A comparison between the old and new recruitment curves shows a considerable reduction of the activation threshold from about 225 V/m (dashed black line) to 30-40 V/m half maximum (red line) when considering the effects of microscopic electric fields.

**Fig. 4:**
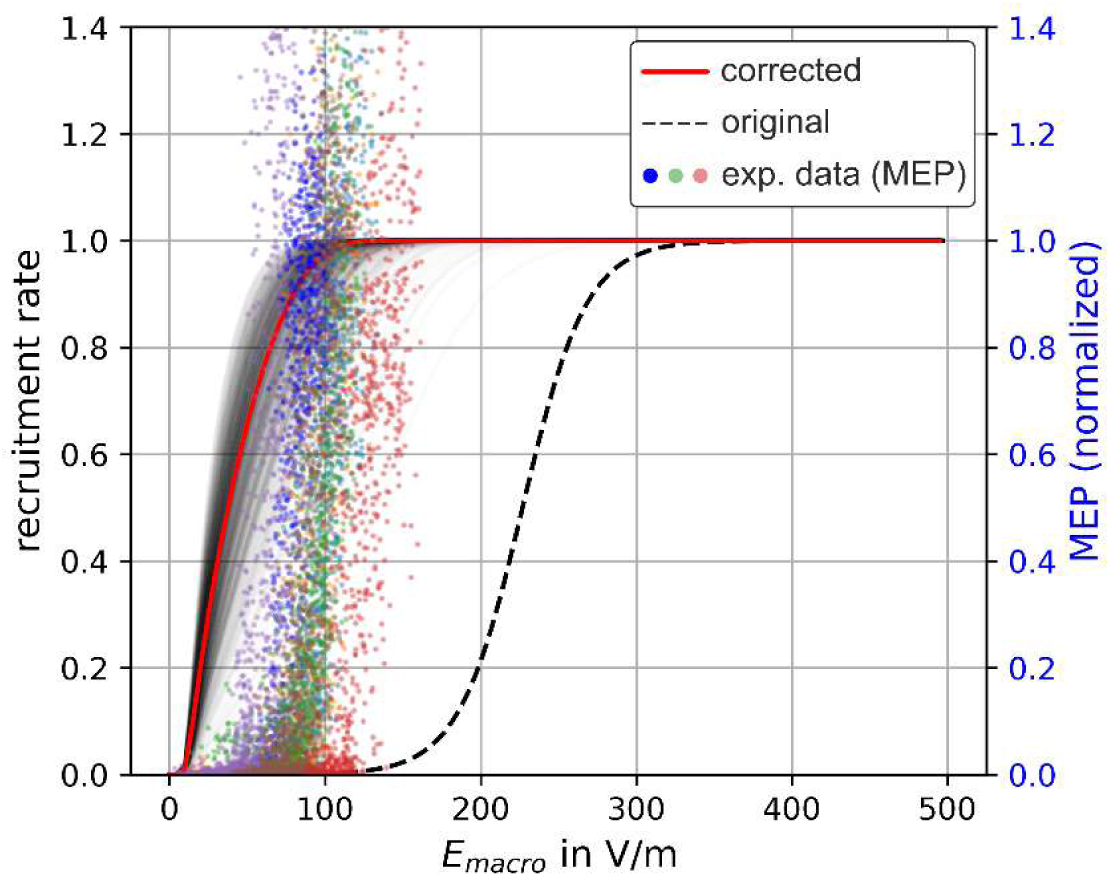
Uncorrected and corrected recruitment rates in comparison to experimental MEPs. Dashed black line: original recruitment rates from Weise et al. (2023b) of L2/3 PC for *θ* = 0° and 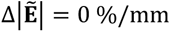 considering homogeneous macroscopic electric fields and a biphasic TMS pulse, without taking into account microscopic electric field effects. Red line: corrected recruitment rates determined from eqs. (1) and (2) using the probability density *p*(*s_e_*) of the electric field scaling factor between microscopic and macroscopic electric fields from Fig. 2, together with the recruitment rate determined using macroscopic electric fields from Weise et al. (2023b) assuming an average number of axon terminals of N=35. Grey lines: Recruitment rate curves after sampling the number of axon terminals N from a normal distribution with the given mean of N=35 and standard deviation of 13.7. Colored dots: MEPs as function of the external macroscopic electric field determined experimentally in Numssen et al. (2021) after motor mapping (different colors represent different subjects).

To enable a comparison of the results with experimental data, we also present the I/O curve of motor evoked potentials (MEPs) of the first dorsal interosseous (FDI) of a representative subject from Numssen et al. (2021) as a function of the macroscopic electric field calculated in that study after successful motor mapping.

A comparison between the recruitment curves shows a clearly improved correspondence to the field strength values observed in the experiment when considering microscopic electric field effects. Note that it is straightforward to apply the correction of the recruitment rates to all neuronal populations, i.e. L2/3 PC, L4 S/N/LBC, and L5 PC presented in Weise et al. (2023b) including different electric field angles with respect to the cortical normal direction (*θ* = [0°, 180°]) and linear field gradients along that direction 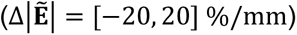. As an example, Fig. 5 shows the dependences of the uncorrected and corrected recruitment rates of L2/3 PC stimulated with biphasic TMS pulses. The correction was applied to all recruitment rate interpolators from all neuronal populations considered in Weise et al. (2023b) and can be downloaded from Weise et al. (2025). Fig. 5 (c) and (d) show the recruitment rate curves for field angles *θ* = 0°, *θ* = 90°, and *θ* = 180° without and with microscopic electric field correction, respectively. It can be observed that the field correction influences the shape of the recruitment rate curves. The directional sensitivity is still present with comparable ratios. Comparing the required e-field values at a recruitment rate of 0.5 for example results in the case of uncorrected macroscopic fields *E*(*θ* = 90°) = 260 V/m, *E*(*θ* = 0°) = 227 V/m, and *E*(*θ* = 180°) = 231 V/m, which results in 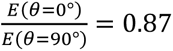 and 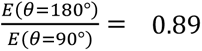. With microscopic field correction, the electric fields are *E*(*θ* = 90°) = 43.3 V/m, *E*(*θ* = 0°) = 37.7 V/m, and *E*(*θ* = 180°) = 39.5 V/m, resulting in 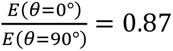 and 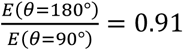.

**Fig. 5:**
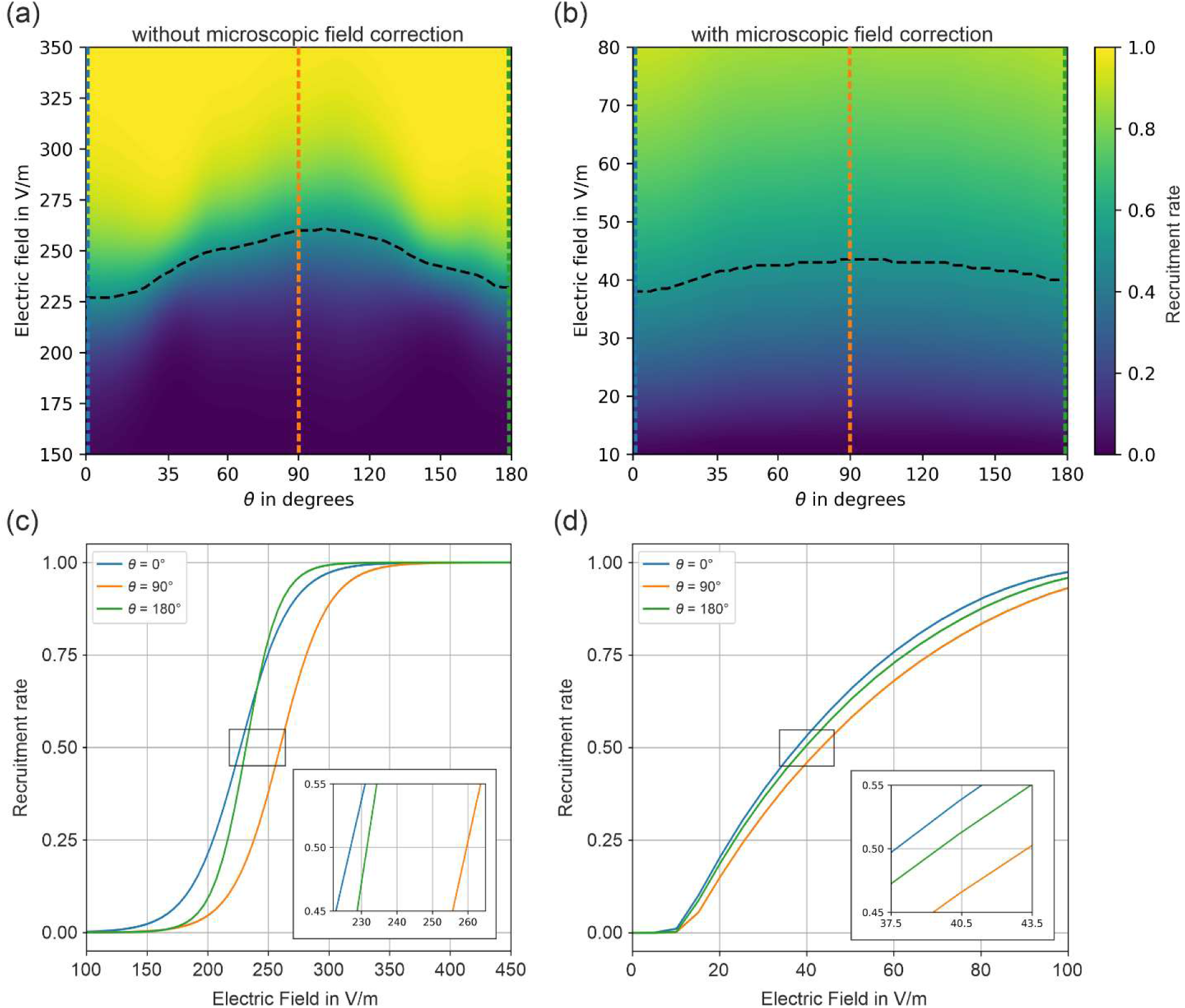
Recruitment rates of L2/3 PC without (a) and with (b) microscopic field corrections stimulated by biphasic TMS pulses for different electric field angles. No e-field gradient 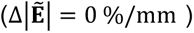 was assumed. (a) The neuronal recruitment rate from Weise et al. (2023b) did not consider microscopic electric field variations, yielding e-field thresholds of above 200 V/m. (b) Recruitment rate of a neuronal population taking electric field variations from Fig. 3 into account yields thresholds of below 50 V/m. (c) and (d) Recruitment rates for field angles of *θ* = 0°, *θ* = 90°, and *θ* = 180° without and with microscopic field correction, respectively.

## 4. Discussion

In recent years, TMS advances have been supported by macroscopic field stimulations tailored to individual head and brain morphology (e.g., Makarov et al., 2020; Thielscher et al., 2015). These models help explain and optimize TMS applications by quantifying cortical stimulation strength in terms of a physical entity: the induced (macroscopic) electric field strength (Caulfield et al., 2021; Numssen et al., 2024). However, using the macroscopic (voxel-based) electric field as a proxy for neural excitation is a simplification, as neurons respond to the electric field in a non-linear fashion. Significant efforts to account for this relationship have been made by detailed mechanistic modeling of neurons at different spatial and temporal scales, from simulated single-cell responses to single pulses (Aberra et al., 2020; Weise et al., 2023b) to network plasticity after repetitive TMS (rTMS; Shirinpour et al., 2021). However, these approaches have not converged on the neuronal activation threshold. Single biphasic pulse TMS of the motor cortex typically yields MEP thresholds of 60-100 V/m macroscopic electric fields (Numssen et al., 2021; Numssen, Kuhnke et al., 2024; Rosanova et al., 2009; Caulfield et al., 2024), while EEG recordings detect changes in cortical activity for rTMS at around 35 V/m (Zmeykina et al., 2020). This variability across methods may reflect different noise levels, which affects detectability. In contrast, single-cell simulations of macroscopic electric fields have yielded threshold estimates above 200 V/m for single monophasic pulses (Aberra et al., 2020; Weise et al., 2023b).

In this study, we investigated local electric field perturbations when considering neural structures at the microscopic level in the electric field simulations. This enabled us to bridge the gap between activation thresholds observed in experiments with respect to macroscopically computed electric fields and those predicted by detailed microscopic simulations of neurons. Our approach lowers this threshold estimate to about 30…40 V/m (Fig. 4), which is in the order of EEG experiments (see above). Also, the corrected half-maximum values around 50 V/m are much closer to the values seen for experimental data using recorded MEPs together with macroscopic field calculations (e.g., 60 V/m in Numssen, Kuhnke et al., 2024). Hence, the recruitment rates computed using the more realistic (i.e., microscopic) electric fields provide a reasonable proxy for the activation thresholds of hand muscles (and, thus, MEPs). This was not the case when using the macroscopically estimated field strength as the field at the neuronal membranes (see Fig. 4).

Note, however, that our model describes the recruitment rate of neurons in the motor cortex, which is not identical to muscle activation reflected by MEP, which involves further downstream processing. Thus, including these processes (e.g., cortical dynamics, long range axonal transmission, spinal dynamics, and muscle fiber activation function) is expected to yield an even more accurate prediction of the MEP.

One key precondition to our approach is the ability to predict microscopic fields at very fine detail by means of large-scale numerical computations. To this end, we utilized results from a method using the BEM-FMM to model perturbations of an impressed electric field within a microscopically realistic brain tissue sample, with many tightly spaced neuronal cells and other structures (Qi et al., 2025). The obtained results (Fig. 1 and 2) demonstrate strong local field perturbations due to the presence of membranes. The derived probability density function of the ratio between microscopic and macroscopic electric fields allowed us to apply a statistical correction to the previously determined recruitment rates by Weise et al. (2023b), who assumed equality between the macroscopic field and the local field at the membranes. Critically, this approach entails that the time consuming numerical field computations have to be performed only once (for a particular type of tissue) and are then reused in the form of a statistical distribution to correct recruitment rates derived from simple macroscopic field estimations. Note, however, that the probability density function of the microscopic/macroscopic field ratio is potentially tissue dependent. Here, we applied an available sample of mouse visual cortex to the activation threshold in human motor cortex. Although we assume that the overall trend in microscopic field distortion would be similar in all cortical tissues, we certainly lose some accuracy. Potentially, cell morphologies, cell alignment (anisotropy), cell density, and other factors exercise an influence on the scaling factors. For the future, we envision the construction of a library of different tissue types, for each of which a microscopic field simulation is performed and a separate probability density function is derived. Such research would also shed light on the question to what extent this relationship can be actually generalized and how much we have to take into account local tissue properties.

Another limiting factor of the model refers to the neural density in the sample. In the model of Qi et al. (2025), the neural density is about 30±5%, which is significantly less than previously reported values of about 70% (Pilgrim et al., 1982; Lehmenkühler et al., 1993; Mota et al., 2014). This discrepancy is due to the technical limitation that all neurons with soma outside the sample were omitted and consequently their dendrites and axons are also missing in the segmentation leading to a decrease in neural density. This is expected to have an influence on the electric field scaling distribution. In order to reduce this influence, larger samples in the millimeter scale would be necessary.

The simulations by Weise et al. (2023b) provide detailed insight into the statistics of the activation thresholds for various neuronal populations, and how they depend on parameters of the external electric field, such as the angle of incidence and field gradient. Already these simulations were complex and computationally expensive, to a degree that impedes usage in whole-head models. Here, we characterized the microscopic electric field considering the mutual influence of the neurons in a realistic cubic sample of mouse visual cortex, based on simulations by Qi et al., 2025, which were even more time-consuming. Theoretically, all simulations carried out by Weise et al. (2023b) would also have to consider microscopic field effects. However, this would lead to an exponential increase in the computing time and is currently far from being feasible. It would also require a much larger tissue sample, because in the one currently utilized, large portions of the neuronal arbors were cut out, rendering direct threshold simulations biased (see Qi et al., 2025). The approach presented here elegantly decouples both problems and considers the determination of the thresholds in the homogeneous field for different cell types and the deviation of microscopic from macroscopic fields separately. Both approaches are then combined by statistically incorporating the electric field deviations together with the recruitment curve in eqs. (1) and (2). Currently, this is the computationally tractable approach to quantify the impact of microscopic electric field perturbations on the firing thresholds and recruitment curves caused by TMS. It represents a significant breakthrough by aligning neural modeling with experimental realities for the first time. Apart from a better understanding of the TMS effect, this may open the door to more systematic procedures for the design of effective stimulation protocols (Shaner et al., 2023).

A promising extension to the current approach of estimating the thresholds would be the use of bidomain models (Czerwonky et al. 2023; Fellner et al. 2022) of the neurons in combination with the microscopic field simulations, instead of the one-dimensional cable equation in conjunction with a much larger 1 mm^3^ MICrONS mouse brain sample (MICrONS Consortium 2021), which includes considerably better developed axonal arbors and ∼75,000 neurite cells. However, the necessary computing power would be immense and is currently not yet available.

Another interesting question would be whether and to what extent the same effects would apply for transcranial electric stimulation (TES). As both methods are based on externally caused electric fields (Peterchev et al., 2012), one could speculate that also with TES the microscopic fields at the cell membranes are larger than the macroscopically computed ones. However, the different ways of impressing that field (magnetic induction vs. externally applied voltage) may give rise to differences, which need to be explored in future computational research.

## Data availability statement

The electric field data used in this study is published by Qi et al. (2025) via BossDB (https://bossdb.org/project/makaroff2024). It includes post-processed cell CAD models, microcapillary CAD models, post-processed neuron morphologies, extracellular electric fields and potential distributions. The derived electric field scaling distribution and recruitment rate operators are publicly available in a repository (Weise et al. 2025, https://osf.io/wu4sa/).

## Acknowledgements

This study has received support from BMBF grant 01GQ2201 (KW, TRK), NIMH grant R01MH130490 and NIBIB grant R01EB035484 (SNM) and NINDS grant 1R01NS112996 (MB).

## Credit Authorship Statement

KW, SNM, and TRK formulated the overarching research goals and aims. SNM computed and curated the electric field data. KW and TRK developed the threshold correction approach. KW and SNM prepared the visualizations. KW, SNM, and TRK acquired the financial support. SNM provided the computing resources for the computationally expensive electric field calculations. KW and TRK wrote the initial draft. SNM contributed to the methods and discussion section and critically reviewed the whole manuscript. ON and MB continued to the analysis and discussion, and critically reviewed the manuscript.

